# PARbars: cheap, easy to build ceptometers for continuous measurement of light interception in plant canopies

**DOI:** 10.1101/481218

**Authors:** William T. Salter, Andrew M. Merchant, Matthew E. Gilbert, Thomas N. Buckley

**Affiliations:** School of Life and Environmental Sciences, Sydney Institute of Agriculture, The University of Sydney, Brownlow Hill, NSW, Australia; Matthew E. Gilbert Department of Plant Sciences, University of California, Davis, Davis, CA, USA

**Author notes:** Corresponding author: William T. Salter.

**Keywords:** Canopy, ceptometer, photosynthetically active radiation, plant area index, phenotyping, transmittance

## Abstract

**Short Abstract:** Detailed instructions on how to build, calibrate and collect research quality data from PARbar ceptometers are presented.

**Long Abstract:** Ceptometry is a technique used to measure the transmittance of photosynthetically active radiation through a plant canopy using multiple light sensors connected in parallel on a long bar. Ceptometry is often used to infer properties of canopy structure and light interception, notably leaf area index (LAI) and effective plant area index (PAI_eff_). Due to the high cost of commercially available ceptometers, the number of measurements that can be taken is often limited in space and time. This limits the usefulness of ceptometry for studying genetic variability in light interception, and precludes thorough analysis of, and correction for, biases that can skew measurements depending on the time of day. We developed continuously logging ceptometers (called PARbars) that can be produced for USD $75 each and yield high quality data comparable to commercially available alternatives. Here we provide detailed instruction on how to build and calibrate PARbars, how to deploy them in the field and how to estimate PAI from collected transmittance data. We provide representative results from wheat canopies and discuss further considerations that should be made when using PARbars.

## Introduction

Ceptometers (linear arrays of light sensors) are used to measure the proportion of photosynthetically active radiation (PAR) intercepted by plant canopies. Ceptometers are used widely for agricultural crop research due to the relatively straightforward nature of measurements and simplicity of data interpretation. The basic principle of ceptometry is that transmittance of light to the base of a plant canopy (*τ*) is dependent on the projected area of light absorbing materials above. Measurements of PAR above and below the canopy can therefore be used to estimate canopy traits such as leaf area index (LAI) and effective plant area index (PAI_eff_) (which includes stems, culms and reproductive structures in addition to leaves)^1-3^. Reliability of PAI_eff_ estimates inferred from τ is improved by modelling the effects of the beam fraction of incoming PAR (*f*_b_), the leaf absorptance (*a*) and the effective canopy extinction coefficient (*K*); K in turn depends on both the solar zenith angle (*θ*) and the leaf angle distribution (*χ*)^1,4-6^. It is common practice to correct for these effects. However, there are other biases that have not received due consideration in the past due to methodological and cost limitations.

We recently identified significant time-dependent bias in instantaneous ceptometry measurements of row crops, such as wheat and barley^7^. This bias is caused by an interaction between row planting orientation and solar zenith angle. To overcome this bias, continuously logging ceptometers can be mounted in the field to monitor diurnal cycles of canopy light interception and then daily averages of *τ* and PAI_e__ff_ can be calculated. However, continuous measurements are often unfeasible due to the prohibitively high cost of commercially available ceptometers – often several thousand US dollars for a single instrument – and the requirement for measurements of many field plots. The latter is particularly evident in the ‘-omics’ era where many hundreds of genotypes are required for genomic analyses, such as genome wide association studies (GWAS) and genomic selection (GS) (for review see Huang & Han, 2014^8^). We recognised that there was a need for cost-effective ceptometers that could be produced in large numbers and be used for continuous measurements across many genotypes. As a solution we designed easy-to-build, high-accuracy ceptometers (PARbars) at a cost of USD $75 per unit. PARbars are built using 50 photodiodes that are sensitive only in the PAR waveband (wavelengths 390 – 700 nm), with very little sensitivity outside this range, negating the use of costly filters. The photodiodes are connected in parallel across a 1 m length to produce an integrated differential voltage signal that can be recorded with a datalogger. The circuitry is encased in epoxy for waterproofing and the sensors operate over a large temperature range (−40 to +80°C), allowing the PARbars to be deployed in the field for extended periods of time. With the exception of the photodiodes and a low-temperature- coefficient resistor, all parts required to build a PARbar can be purchased from a hardware store. A full list of required parts and tools is provided in Table 1. Here we present detailed instructions on how to build and use PARbars for estimation of PAI_eff_ and present representative results from wheat canopies.

**Table 1.**
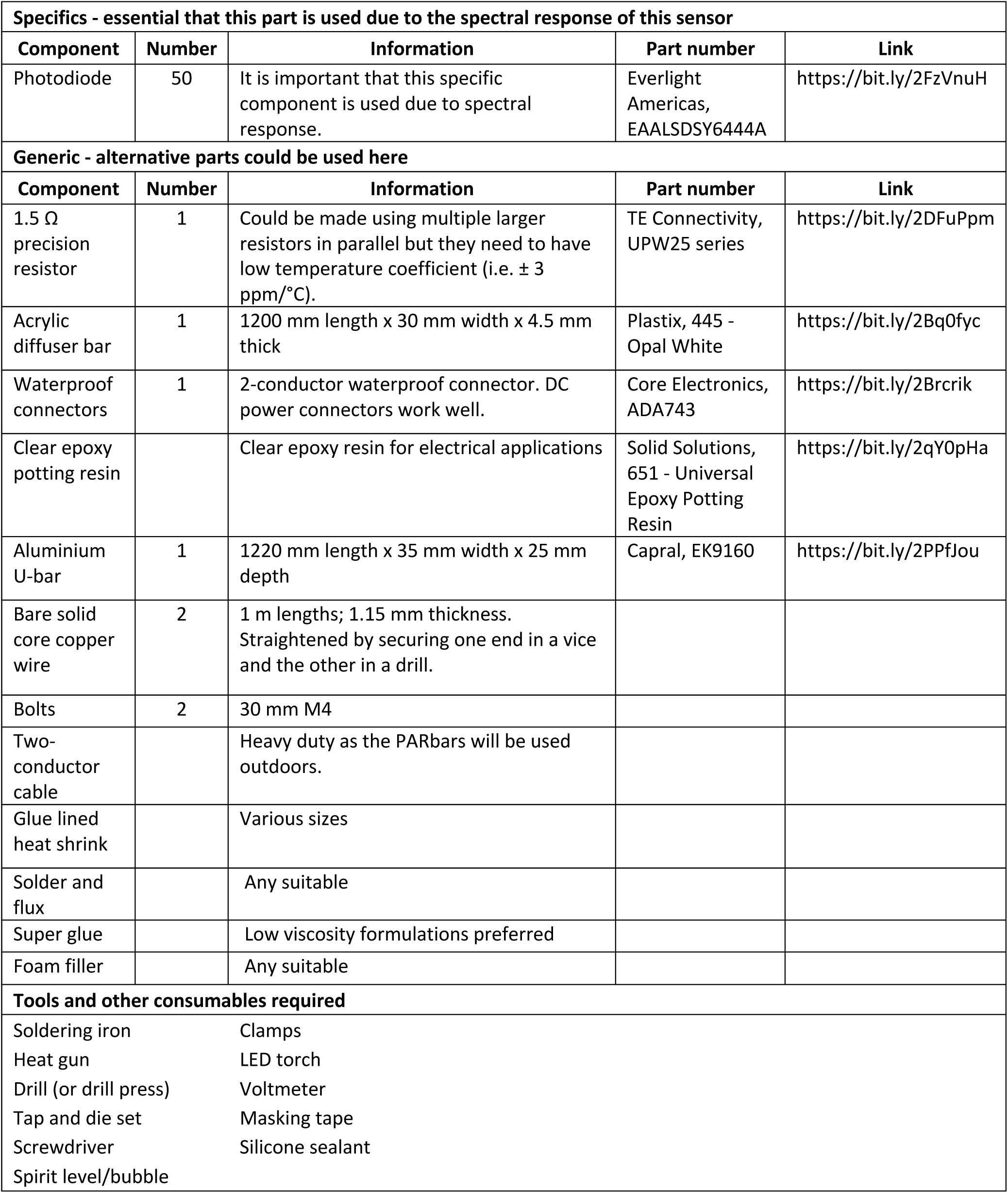
Components and tools required to build a PARbar ceptometer. Note that the photodiode is a specific component, and it is essential that it is used due to its spectral response. All other parts can be obtained from hardware and electronics suppliers, and suitable alternatives to the part numbers stated could be used.

## Protocol

### 1. Building and calibrating the PARbars

1.1) Gather all parts and tools required for assembly in a clean workspace. Note that PARbars can also be produced in batches due to long curing times required at certain points. Note that schematics of a PARbar can be found in Figure 1 for reference.

**Figure 1.**
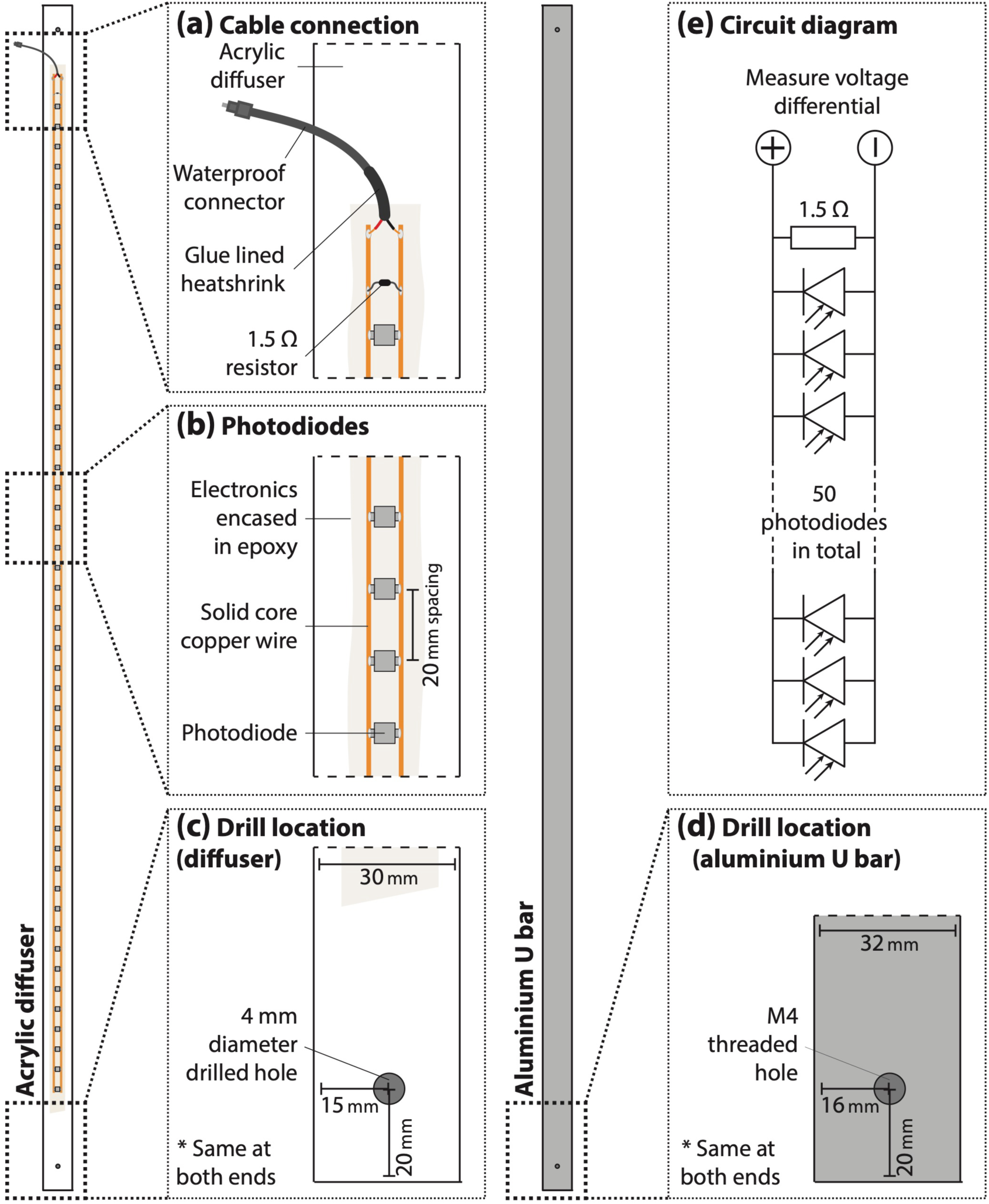
Schematics for the PARbar build. (a) highlights the location and arrangement of the waterproof connector and the internal shunt resistor; (b) highlights the arrangement and spacing of the photodiodes; (c) highlights the drilling locations on the acrylic diffuser bar; (d) highlights the drilling locations on the aluminium U-bar; and (e) shows an electronic circuit diagram of a PARbar.

1.2) Drill a 4 mm diameter hole 20 mm from each end of an acrylic diffuser bar (1200 mm length x 30 mm width x 4.5 mm thickness; 445 – Opal White; Plastix Australia Pty. Ltd., Arncliffe, NSW, Australia). Drill and tap threaded holes in a section of aluminium U-bar to secure diffuser, 20 mm from each end. Drill and tap threaded holes to suit mounting hardware (e.g., a tripod mounting plate).

1.3) Generally, bare copper wire comes on a roll and needs to be straightened before it can be used in the PARbar circuit. Secure one end of a 1.25 m length of wire (1.25 mm diameter) into a vice or clamp and tighten the other end into the grips of a hand drill. Turn on the drill to straighten the wire. Repeat with a second 1.25 m length of bare copper wire.

1.4) Mark the intended locations of the copper wire and the photodiodes along the edge of the diffuser using a fine-tip permanent marker (full schematics can be found in Figure 1).

1.5) Superglue one of the straightened copper wires to the diffuser. Super glue 50 photodiodes (EAALSDSY6444AO; Everlight Americas Inc., Carrollton, Texas) face-down along the diffuser at 20 mm intervals, ensuring that they are in the centre of the diffuser and that all are arranged all in the same orientation such that the large tab sits on the copper wire. Super glue the other copper wire to the diffuser, such that it sits underneath the smaller tabs of the photodiodes.

1.6) Apply some solder flux to the photodiode tabs and solder the photodiodes to the copper wires. Test solder connections by shining a light onto each photodiode individually and checking for a voltage signal across the wires using a multimeter.

1.7) Solder a 1.5 Ω resistor in parallel across the copper wires, this will produce a linear quantum response (*this step is optional, if resistor is not soldered into the PARbar, it can instead be connected in parallel with the PARbar signal inputs on the datalogger*). Low temperature coefficient precision resistors should be used to prevent ambient temperature from influencing the voltage signal at a given light level.

1.8) Solder the male end of a waterproof DC connector (ADA743; Core Electronics, Adamstown, NSW, Australia) to the ends of the copper wire and seal the connections using glue lined heat shrink tubing.

1.9) Using silicone sealant, create a continuous silicone barrier around the circuity to form a fluid-tight well. Once the sealant has cured, fill the well with epoxy resin (651 – Universal Epoxy Potting Resin; Solid Solutions, East Bentleigh, VIC, Australia).

1.10) When the epoxy resin has hardened (overnight), remove the silicone sealant using a razor blade. Bolt the diffuser to the pre-threaded aluminium U-bar using M4 bolts.

1.11) Use masking tape to secure the diffuser to the aluminium along its whole length and fill the space inside the ceptometer with polyurethane foam filler. Once the foam filler has set (overnight), remove the masking tape. The ceptometer is now complete.

1.12) Solder the female end of the DC connector to a length of two-conductor cable, which will be connected to the datalogger, and seal the connections with glue lined heat shrink.

1.13) The PARbar should be calibrated against a quantum sensor (such as LI-190R; LI-COR, Lincoln, Nebraska, USA). Connect both sensors to a datalogger (such as CR5000; Campbell Scientific, Logan, Utah, USA) and set them outside in full sun on a level plane (level with a spirit level or spirit bubble). Log the outputs of both sensors for a full diurnal cycle. Plot a calibration curve (such as Figure 2) to convert the raw voltage signal from the PARbars to PAR using the quantum sensor output.

**Figure 2.**
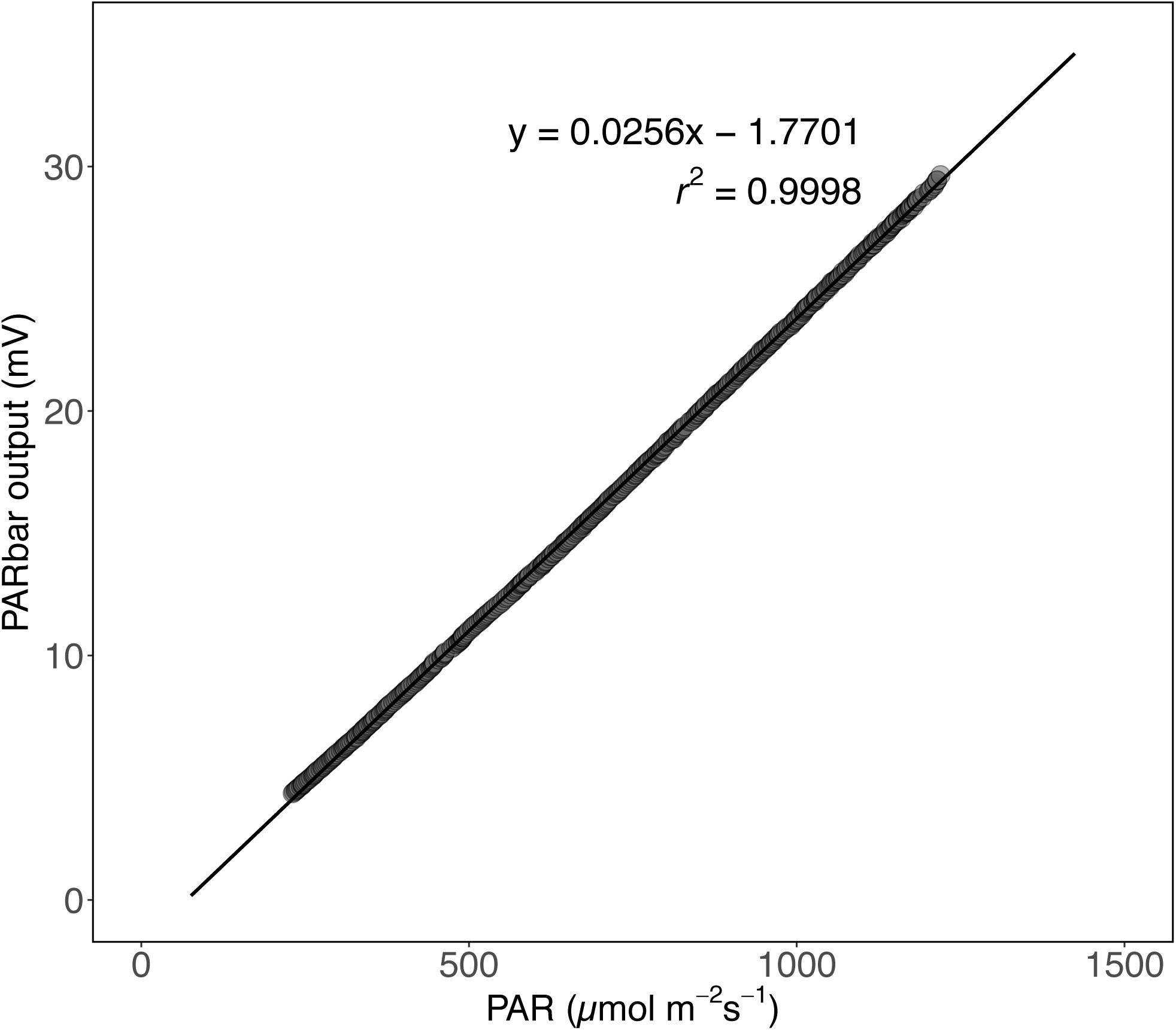
A representative PARbar calibration curve, showing the relationship between the differential voltage output of a PARbar and the photosynthetic photon flux density from a LI-COR LI-190R quantum sensor.

### 2. Installation in the field

2.1) To infer PAI_eff_, one PARbar (or quantum sensor) should be set up above the canopy with the other PARbars inserted below the canopy at a 45° angle to row planting. The PARbar above the canopy can be mounted on a tripod. All PARbars should be levelled using a spirit level or bubble. It is strongly encouraged that data is sampled across a full diurnal cycle due to time dependent bias of instantaneous measurements^7^.

2.2) Connect the PARbars to a datalogger using cables made in step 1.11 and commence logging at desired sampling interval. Remember to connect each in parallel with a 1.5 Ω low temperature coefficient precision shunt resistor if this was not integrated into the PARbar design.

2.3) Collect data from the datalogger and transfer to a computer. Differential voltage data can be converted to PAR using the calibration for each PARbar.

### 3. Calculation of effective plant area index (PAI_eff_)

3.1) PAI_eff_ can be calculated for each time point in the dataset using the following equations (provided in the manual for the AccuPAR LP-80 ceptometer; Decagon Inc., Pullman, WA, USA^6^):

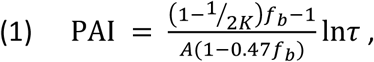

where *A* = 0.283 + 0.0785*a* – 0.159*a*^2^ (in which *a* is leaf absorbtance), *τ* is the ratio of below- to above-canopy PAR, and *K* and *f*_*b*_ are modelled by Equation 2^4^ and Equation 3^9^, respectively:

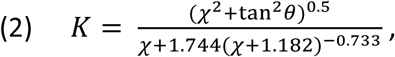

where *χ* is a dimensionless parameter describing leaf angle distribution, *θ* is the solar zenith angle, and

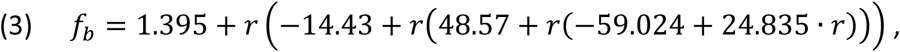

where *r* is PAR above the canopy (PAR_above_) as a fraction of its maximum possible value (PAR_above,max_ = 2550·cos*θ*); i.e. *r* = PAR_above_/PAR_above,max_. For wheat we assumed *a* = 0.9 and *χ* = 0.96 (the latter value was given for wheat by Campbell and van Evert (1994)^10^). An R script is provided as a supplementary file for automated processing of large datasets.

### Representative results

A representative calibration curve for a PARbar is shown in Figure 2. The differential voltage output of a PARbar is linearly proportional to the PAR output from a quantum sensor, with *R*^2^ = 0.9998. PARbars were deployed in wheat canopies and logged every 20 s across the development of the plants. A typical diurnal timecourse of the canopy light environment collected using a PARbar on a clear sunny day is shown in Figure 3 (raw transmittance data and corrected PAI are shown for comparison). Figures 3b and 3c demonstrate the bias that could be introduced by taking instantaneous ceptometry measurements at various times of day (as per Salter *et al*. 2018^7^). The wheat plots used for the collection of this data had a row planting orientation due north-south with transmission of light to the lower canopy peaking at 12:30 (Figure 3b). If an instantaneous measurement were to be taken at this point, PAI would be underestimated whilst if it was taken in the morning or afternoon it may be overestimated. The weatherproof PARbars can also be deployed in the field for longer time periods; Figure 4 demonstrates how the PARbars could be used to monitor how canopy light environment changes as the plants develop.

**Figure 3.**
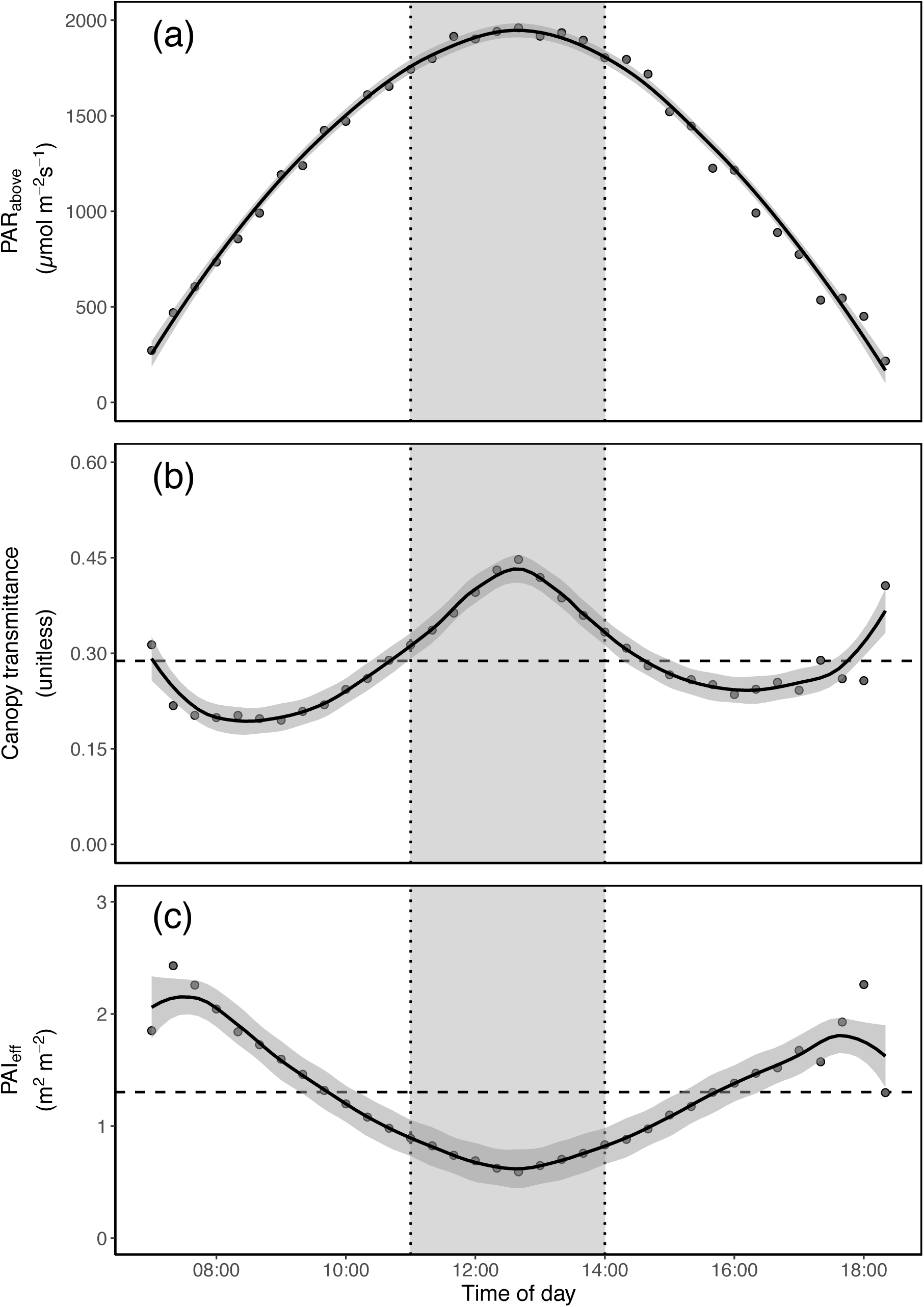
Representative daily timecourse data collected on a clear day using PARbars in wheat canopies at anthesis in Canberra, Australia (−35°12’00.1008", 149°05’17.0988”). (a) shows the PAR measured above the canopy, (b) the uncorrected transmittance data (i.e. PAR_above_/PAR_below_), and (c) the effective plant area index (PAI_eff_), corrected for the beam fraction of incoming PAR (*f*b), the leaf absorptance (*a*) and canopy extinction coefficient (*K*). Data points shown in (b) and (c) are means (*n* = 30), solid lines are LOESS local regressions fitted in R (*a* = 0.5), shaded areas are standard errors of the fit and the dashed horizontal lines represent the daily means. The shaded area between the dotted lines is the time window (1100 – 1400h) recommended for instantaneous ceptometer measurements in wheat by CIMMYT^11^.

**Figure 4.**
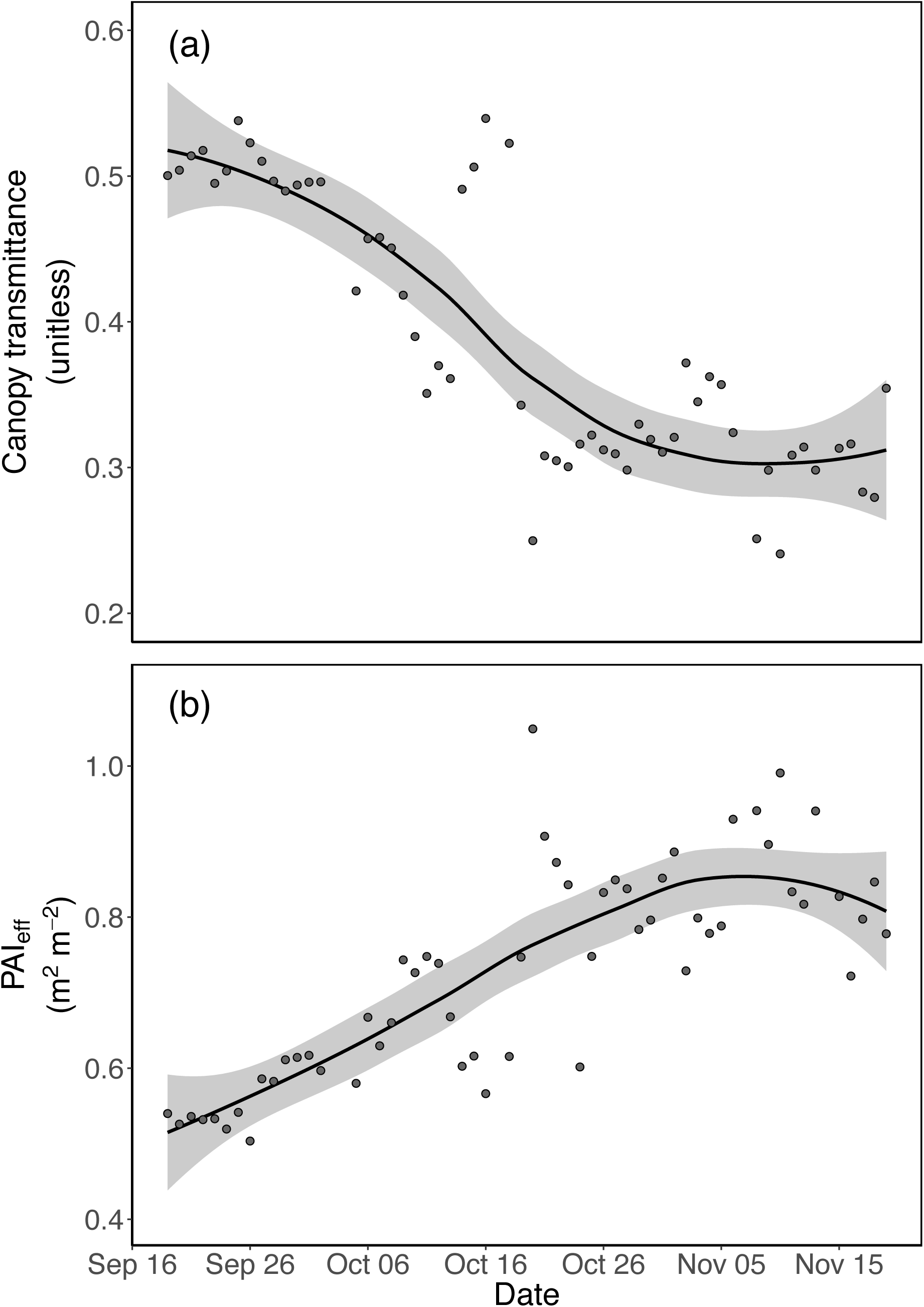
Representative data collected across a growing season (from early tillering to anthesis) using PARbars deployed in wheat canopies in Canberra, Australia (−35°12’00.1008", 149°05’17.0988”). (a) shows the uncorrected transmittance data and (b) the effective plant area index, corrected for the beam fraction of incoming PAR (*f*b), the leaf absorptance (*a*) and canopy extinction coefficient (*K*). Data points shown represent daily means for the period 1000 – 1400h (*n* = 30). Solid lines are LOESS local regressions fitted in R (*a* = 0.75), shaded areas are standard errors of the fit. Raw data was not included in further analysis if PAR_above_ was < 1500 µmol m^-2^ s^-1^ and if PAR_below_/PAR_above_ was > 1.

## Discussion

The quality of data collected with PARbars make them an alternative to expensive commercial ceptometers, yielding an *R*^2^ > 0.99 when calibrated against a LI-COR Li-190R quantum sensor (Figure 2). Similar high correlations were found for 68 PARbars used in previous work^7^. As with most commercial light sensors, calibrations differ among PARbars so their output must be converted using their specific individual calibrations. Recently, there has been a growing interest in novel high-throughput plant phenotyping technologies for the estimation of canopy traits (for review see Yang *et al*., 2017^12^). Whilst these methods are promising in that they produce huge amounts of data they are typically very indirect and require validation against conventional techniques. PARbars could serve as a cost-effective, ground-based validation tool for these new techniques.

Our previous work^7^ highlighted that continuous ceptometry measurements across diurnal cycles are required for reliable estimation of PAI of row crop canopies, due to interactions between solar zenith and row-planting orientation that could bias instantaneous measurements. This can also be seen in Figure 3. The low production cost of PARbars make them a viable option for continuous measurements in the field. With sufficient numbers of PARbars, continuous measurements can be performed in all field plots. Alternatively, PARbars can provide continuous measurements in just a few plots in order to characterize row- orientation biases to develop time-specific correction functions for instantaneous measurements (for more information see Salter et al. 2018^7^). Another key benefit of using continuous ceptometry is the ability to capture short fluctuations in *τ* over time (sunflecks and shadeflecks), caused by clouds passing overhead, movement of the canopy, etc. Photosynthesis is known to be highly sensitive to small changes in environmental conditions and ‘dynamic’ changes in photosynthesis are now thought to be important in driving crop yield (for review see Murchie *et al.*, 2018^13^). PARbars installed in the field with a suitably short logging interval could be used to capture these short fluctuations and provide better understanding of the dynamic nature of plant canopies.

PARbars can also be installed in the field for extended periods of time, as shown in the example in Figure 4. One notable exclusion from the current PARbar design that could be considered for long-term monitoring is a means to distinguish between direct beam and diffuse components of incoming PAR above the canopy. As diffuse radiation penetrates deeper into the canopy than direct sunlight^14^, transmittance will be increased and PAI_eff_ will be underestimated. When all radiation is diffuse, PAI is directly proportional to the logarithm of 1/*τ* rather than the relationship shown in Equation 1^15^. The lack of a diffuse component in the data processing steps used in this study may explain some of the day-to-day variation in the data shown in Figure 4. It is possible to find diffuse/direct radiation data in some open access datasets but due to the localised nature of environmental variables that influence incoming PAR (clouds, air pollution, etc.) they are often not applicable to ceptometry data. Cruse *et al*. (2015)^16^ noted that currently available commercial instruments that can measure direct and diffuse PAR are expensive and require regular maintenance, so they designed a simple and cheap apparatus to address this issue. Their system consists of a quantum sensor that is routinely shaded by a motorised, moving shadowband and allows for continuous measurement of total, direct and diffuse PAR. The sensor used in the Cruse *et al*.^16^ system could be replaced with the same photodiode used in PARbars to further reduce cost, and may be easily incorporated into the existing PARbar setup. These measurements could be integrated into the data processing pipeline and would further enhance reliability of estimates of PAI_eff_.

The PARbars that we present in this paper were designed specifically for use in row crops, such as wheat and barley, but the handmade design could easily be modified for a user’s specific requirements. For example, the shunt resistor could be changed to provide linearity at lower PAR ranges, or for versatility a low-temperature coefficient precision potentiometer could be used to change the linear range as necessary. The photodiodes could also be used individually as quantum sensors, allowing the user to capture spatial as well as temporal variation within individual canopies for a much lower cost than would have been possible previously. This could be of particular importance given the growing focus on dynamic photosynthesis under fluctuating light conditions.

Although we used a conventional (and expensive) datalogger for the data presented in this study, there is scope for dataloggers to also be built using off-the-shelf componentry, enabling the creation of a combined ceptometry and datalogger system on a limited budget. The popularity of so-called ‘maker’ platforms, such as Arduino and Raspberry Pi, offer great promise in this area and one might consider the use of the open-source Arduino-based Cave Pearl project^17^ as a starter for further development. The Cave Pearl dataloggers were designed for environmental monitoring of cave ecosystems so ruggedness and low power demand were key considerations in their design. Similar considerations are relevant for implementation to plant phenotyping work. The cost of the components required for the Cave Pearl datalogger is less than USD $50 per unit and due to the small size of the circuit boards used in this project, datalogging could be directly incorporated into future design of PARbars.

PARbars provide a cost-effective and high-accuracy alternative to commercially available ceptometers. They do not require specialist expertise to build, nor for the interpretation of resulting data. Consequently, PARbars could be widely adopted in the plant phenotyping community – including by those who generally use expensive light sensing tools and those who have been unable to access such technology due to budget restrictions. The ‘do-it-yourself’ nature of PARbars means that they could be adapted to a user’s specific needs with added flexibility for future development and adaptation of this technology for a range of purposes.

## Supporting information

## Acknowledgements

The authors would like to thank Dr. Richard Richards and Dr. Shek Hossain at CSIRO, Agriculture and Food for access to and management of the field plots used for this research. This research was supported by the International Wheat Yield Partnership, through a grant provided by the Grains Research and Development Corporation (US00082). TNB was supported by the Australian Research Council (DP150103863 and LP130100183) and the National Science Foundation (Award #1557906). This work was supported by the USDA National Institute of Food and Agriculture, Hatch project 1016439.

