## Supplementary material for "PARbars: cheap, easy to build ceptometers for continuous measurement of light interception in plant canopies"

```r
# R script for calculation of PAI from PARbar ceptometry data

# ------Input data file conditions------
# .csv data file "PARbardata.csv" should have columns containing:
# - Column 1 - Date and time (titled "Dateandtime; format "yyyy-mm-dd hh:mm:ss")
# - Column 2 - Latitude (titled "Latitude"; decimal format, e.g. )
# - Column 3 - Longitude (titled "Longitude"; decimal format, e.g. )
# - Column 4 - PARabove (titled "PARabove"; should already be calibrated)
# - Column 5 onwards - PARbelow (Titled "PARbar1 - PARbar##"; should be pre-calibrated, i.e. umol
m^-2 s^-1 values, not mV).

# ------Set working directory------
# First manually set your working directory
# setwd(enter directory here)

# ------Set timezone------
# Then set your timezone (to help find your timezone, use this link: http://twiki.org/cgi-
bin/xtra/tzdatepick.html )
Timezone <- "Australia/Sydney"

# ------Load packages------
# Install (if necessary) and load required packages
# install.packages("GeoLight")
library(GeoLight)

# ------Load input data file "PARbardata.csv"------
# Load data into dataframe
RAWPARdata <- read.csv("PARbardata.csv")

# ------Automated processing------
# Processing is automated from here, outputting a .csv file "PAIouput.csv to working directory
# Split input dataframe
PAIdataframe <- RAWPARdata[1:4]
RAWPARdata[1:4]=NULL

# Calculate transmittance for each PARbar
Transmittance <- RAWPARdata/PAIdataframe$PARabove
names(Transmittance) <- paste("Tr", names(Transmittance))

# Compute zenith angles and output to dataframe
sunny <- solar(as.POSIXct(PAIdataframe$Dateandtime,tz= Timezone))
PAIdataframe$Zenithangle <- zenith(sunny,lon=PAIdataframe$Longitude,lat =
PAIdataframe$Latitude)
PAIdataframe$Zenithangle <- (pi*PAIdataframe$Zenithangle)/180

# Set leaf angle distribution (chi) and leaf absorbtivity (abs), and calculate bigA
chi <- 0.96
abs <- 0.9
bigA <- 0.283+0.785*abs-0.159*abs^2

# Calculate PARabovemax
```

```
PAIdataframe$PARabovemax <- 2550*cos(PAIdataframe$Zenithangle)

# Calculate ratioPARabovePARabovemax
PAIdataframe$PARratio <- PAIdataframe$PARabove/PAIdataframe$PARabovemax

# Calculate fb
PAIdataframe$fb <- 1.395+PAIdataframe$PARratio*(-
14.43+PAIdataframe$PARratio*(48.57+PAIdataframe$PARratio*(-
59.024+24.835*PAIdataframe$PARratio)))

# Calculate K
PAIdataframe$K <- ((chi^2 +tan(PAIdataframe$Zenithangle)^2)^0.5)/(chi+1.744*(chi+1.182)^-0.733)

# Calculate PAI, output to new df
PAIvalues <- (((1-0.5*PAIdataframe$K)*PAIdataframe$fb-1)/(bigA*(1-
0.47*PAIdataframe$fb)))*log(Transmittance)
names(PAIvalues) <- paste("PAI", names(PAIvalues))

PAIoutput <- cbind(PAIdataframe[1:4],Transmittance,PAIvalues)

# Combine dataframes and output to csv file
PAIoutput <- cbind(PAIdataframe[1:4],Transmittance,PAIvalues)
write.csv(PAIoutput,"PAIoutput.csv")

# ------Time interval means------
# Calculate desired time interval means.
# Time interval format "## day","## hours","## min". Example shown is for hourly means.
PAIoutput$Dateandtimegroup <- as.POSIXct(PAIoutput$Dateandtime,tz=Timezone)
PAIoutput$Hour <- cut(PAIoutput$Dateandtimegroup, breaks = "1 hour")
# Transmittance means
HourlyTRmeans <- aggregate(PAIbigframe[6:33],by=list(PAIbigframe$Hour), mean)
# PAI means
HourlyPAImeans <- aggregate(PAIbigframe[39:65],by=list(PAIbigframe$Hour), mean)
```
